# MORPHIOUS: A Machine Learning Workflow to Naively Detect the Activation of Microglia and Astrocytes

**DOI:** 10.1101/2020.08.17.251843

**Authors:** Joseph Silburt, Isabelle Aubert

## Abstract

In cases of brain injury, degeneration and repair, defining microglia and astrocytic activation using cellular markers alone remains a challenging task. We developed MORPHIOUS, an unsupervised machine learning workflow that utilizes a one-class support vector machine to segment clusters of activated glia by only referencing examples of non-activated glia. Here, glial activation was triggered using focused ultrasound to permeabilize the hippocampal blood-brain barrier. Analyzing the hippocampal sections seven days later, MORPHIOUS identified two classes of microglia which showed characteristic activation features, including increases in ionized calcium-binding adapter molecule 1 expression, soma size, and de-ramification. MORPHIOUS was further used to identify clusters of activated astrocytes, which showed increased expression of glial fibrillary acidic protein and branching. Thus, by only referencing untreated glia morphologies, MORPHIOUS can identify diverse and novel manifestations of glial activation. This provides significant improvements for characterizing glial activation in cases of injury, neurodegeneration, and regeneration.

## Introduction

Ubiquitous across neurodegenerative diseases, microglia and astrocytes represent important glial cell populations which are activated in response to pathology. Depending on the context, this activation can either ameliorate or exacerbate disease progression^1,2^. Importantly, the activation of microglia and astrocytes is accompanied by distinct morphological characteristics. To better understand glial activation, several machine learning approaches have been developed to classify activated states based on their morphology. Commonly, these methods deploy unsupervised learning algorithms (e.g., K-means clustering, hierarchical clustering)^3–7^. In general, these approaches aim to group activated and non-activated glial morphologies into distinct groups based on the similarities of their features. However, given that the activation of microglia and astrocytes exists along a gradient of morphologies^4,7–9^, it remains difficult to define a strict classification boundary to accurately identify activated cells.

Supervised learning algorithms have shown promises in classifying cell types. Supervised learning classifiers learn rules based on patterns in labelled data in order to discriminate between multiple classes of data ^10^. Among many applications, supervised learning algorithms have been used to identify activated microglia following traumatic brain injury^11^, and to distinguish between macrophage activation states ^12^. While powerful, supervised learning classifiers must be provided with labelled data, where the classification of each data point is already known. For many clinical, preclinical, and basic biological problems, including for detecting activated microglia and astrocytes, such datasets are not widely available and must be generated using expert knowledge. Moreover, because supervised classifiers are trained on predefined classes, supervised classifiers suffer from an inability to discover new classifications, which is of interest to biologists^10^.

We developed a method to identify activated astrocytes and microglia using a one-class support vector machine. Support vector machines in general have been widely used in biology and are both capable of modelling significant complexity, while also regularizing against overfitting ^13,14^. Traditionally, support vector machines are supervised, and determine a decision boundary by evaluating the largest margin from which to separate classes of data. By contrast, one-class support vector machines require the input of a single baseline class and the selection of a probability quantity (i.e., nu), which is proportional to the probability that a new observation outside the decision boundary should be considered normal ^15^. Thus, data can be classified based on patterns learned purely from a baseline class.

Using a one-class support vector machine, we developed the MORPHological Identification of Outlier clUSters (MORPHIOUS) procedure, a novel approach to identify classes of glial activation. MORPHIOUS learns the feature patterns of normal, non-activated glial cells, and uses this information to segment regions of glia which are classified to be abnormal, and therefore activated. Our approach employs a naïve definition of what defines glial activation, and therefore facilitates the identification of a robust range of morphologies consistent with a broad definition of activation. In this work we used to quantify the activation of glial cells in the hippocampus of C57bl/6 mice treated with focused ultrasound (FUS) combined with intravenously injected microbubbles, a technology which facilitates the temporary permeabilization of the blood-brain-barrier, and is known to transiently activate microglia and astrocytes^16^. Through our analysis, we show that MORPHIOUS can isolated regions of activated microglia and astrocytes from surrounding non-activated tissue and can be used to identify novel glial activations states based on morphology alone.

## Results

### MORPHIOUS workflow

#### Feature collection

The activation of microglia and astrocytes was induced in mice using a unilateral treatment of focused ultrasound (FUS) in the presence of microbubbles, to the left hippocampus in 14-week old C57bl/6 mice. Mice were sacrificed at 7D post FUS, a timepoint where the activation of both microglia and astrocytes has been previously detected^17^, and processed for immunohistochemical analysis. Microglia were stained with ionized calcium-binding adapter molecule (Iba1), which labels microglial processes and is upregulated with microglial activation^18^. Astrocytes were double-stained with S100 calcium-binding protein beta (S100b) and glial fibrillary acidic protein (Gfap). Gfap, which is upregulated with astrocytic activation^2^, was used to evaluate branching and intensity metrics and S100b was used to count cells and quantify soma characteristics. Using custom ImageJ scripts, features were extracted from whole hippocampal slices by applying a sliding window (Fig. 1-A1, B1). We collected features related to the fluorescence intensity, cellular surface area, branching complexity, cell location, and cell soma shape. Averaged features for each sliding window were extracted. Extracted features were normalized, and principle component analysis (PCA) transformed.

**Fig. 1.**
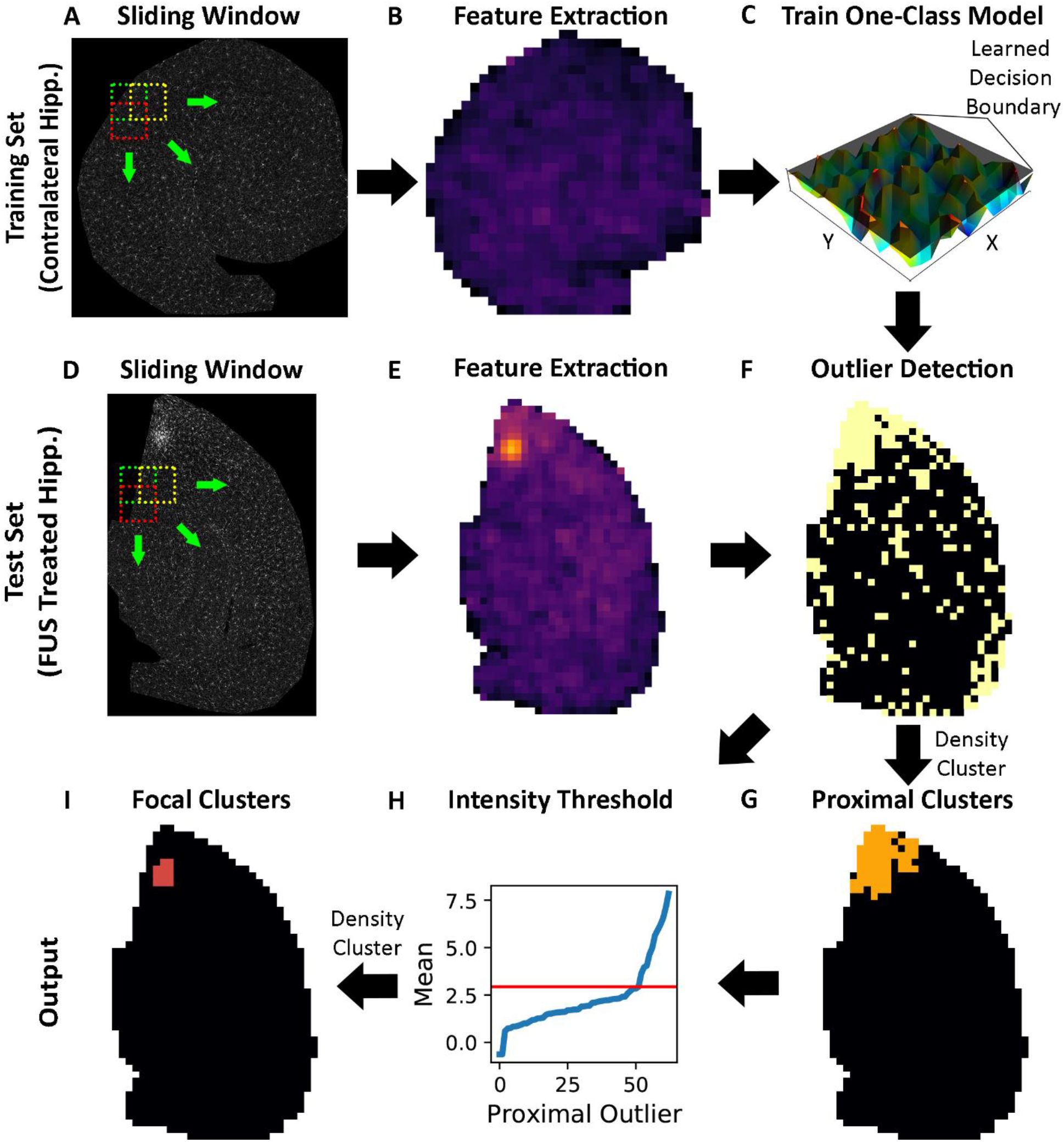
The MORPHIOUS workflow trains a one-class support vector machine to identify activated glial cells. (A) A sliding window is applied contralateral hippocampal sections to extract features. (B) Extracted features are used to generate a spatial map of features. (C) Features from contralateral hippocampal sections are used to train a one-class support vector machine which generates a decision boundary for defining non-activated glia. (D, E) A sliding window is further used to extract features from FUS treated hippocampi. (F) The trained model is applied to FUS treated hippocampi to identify outlier windows. (G) Outliers are spatially clustered using the density-based spatial clustering of applications with noise algorithm (DBSCAN) to identify proximal clusters. (H) To identify focal clusters, the mean intensities of proximal cluster regions are sorted in ascending order, and the elbow point of this curve (red line) is used as a threshold value. (I) DBSCAN is applied windows with a mean intenstiy above this threshold value to identify focal clusters.

#### Automated selection of microglia and astrocytes

To aid in feature collection, we developed two simple protocols using the FIJI morpholibJ package^19^ to automatically count Iba1^+^ and S100b^+^ cell bodies–representing microglia and astrocytes, respectively, and measure cell soma related features. These protocols strongly correlated with manual counts (R^2^: 0.964 for microglia, R^2^: 0.959 for astrocytes) (Supplementary Fig. 1).

#### Building a naïve one-class classifier

We trained a one-class support vector machine using contralateral hippocampal sections for all mice (Fig. 1 A-C). This classifier was subsequently tested on each ipsilateral hippocampal section (Fig. 1 D-E). From this, a putative spatial array of outliers was identified (Fig. 1F). A key insight of our approach is applying the assumption that glial activation occurs in spatial clusters, which are located proximal to stimuli, here specifically due to the FUS treatment. This has been shown to occur in pathologies such as an ischemic vessel^20,21^, or an amyloid-beta plaque^16^. As such, the Density-Based Spatial Clustering of Applications with Noise (DBSCAN) algorithm was used on spatial coordinates of identified outliers, in order to detect consistent clusters of activated/outlier glia (Fig. 1G). These are termed proximal activation clusters. With regards to microglia, we further classified ‘focal’ activation clusters which represent microglia with the most prominent activation features. To delineate the boundary of focal clusters, DBSCAN was applied to proximal cluster windows with a mean Iba1 intensity above a threshold value (Fig. 1I). To calculate this threshold, the mean Iba1 intensity of each proximal outlier was sorted in ascending order and the elbow point of the ensuing curve was used as the threshold value (Fig. 1H, red line).

#### Parameter Tuning

To ensure that identified clusters represent morphologically activated cells, our learning objective was to predict no false-positive. Thus, we applied 10-fold cross validation across all contralateral hippocampal sections to identify hyperparameters where no activation clusters were observed within any contralateral hippocampal sections (Supplementary Fig. 2-3). Within the set of parameters which ensured no false-positive results, we chose parameters which maximized the amount of activated glia observed in FUS-treated hippocampi.

#### Classifying microglia

Using MORPHIOUS, we identified regions of proximal (Fig. 2E) and focal (Fig. 2F) microglial activation. Our results indicate that focal and proximal microglia exhibited a gradient of cell activation features (Fig. 3). When compared to untreated microglia of the contralateral hippocampus, focal microglia exhibited a 1.8-fold increased Iba1 intensity (P<0.0001, Fig. 3A), a 2-fold increase in surface area (Focal: P<0.0001, Fig. 3B), a 1.4-fold increase in soma size (P<0.0001, Fig. 3C), a 1.4-fold reduction in branch length (Focal: P<0.001, Fig. 3D), a 1.8-fold reduction in the number of branches per cell (P<0.0001, Fig. 3E), and a 1.5-fold reduction in nearest neighbour distance (NND) (P<0.0001, Fig. 3F). Moreover, when compared to proximal microglia, focal microglia also exhibited a 1.3-fold increase in Iba1 intensity (P<0.01, Fig. 3A), a 1.4-fold increase in surface area (P<0.001, Fig. 3B), a 1.2-fold increase in soma size (P<0.0001, Fig. 3C), and a 1.2-fold reduction in NND (P<0.05, Fig. 3F). Similarly, proximal microglia exhibited significant, but less pronounced change in Iba1 intensity (P<0.05, Fig. 3A), surface area (P<0.0001, Fig. 3B), and soma size (P<0.05, Fig. 3C), branch length (P<0.001, Fig. 3D), number of branches (P<0.0001, Fig. 3E), NND (P<0.001, Fig. 3F), when compared to contralateral cells. In addition, for all assessed features, unclustered ‘distal’ microglia, were statistically indistinguishable from untreated microglia (P>0.05), rendering them representative of non-activated microglia in the FUS treated hippocampus. These findings are visualized via principal component analysis where focal and proximal activation clusters occupy distinct regions in feature space while distal microglia overlap with untreated microglia (Supplementary Fig. 4).

**Fig. 2.**
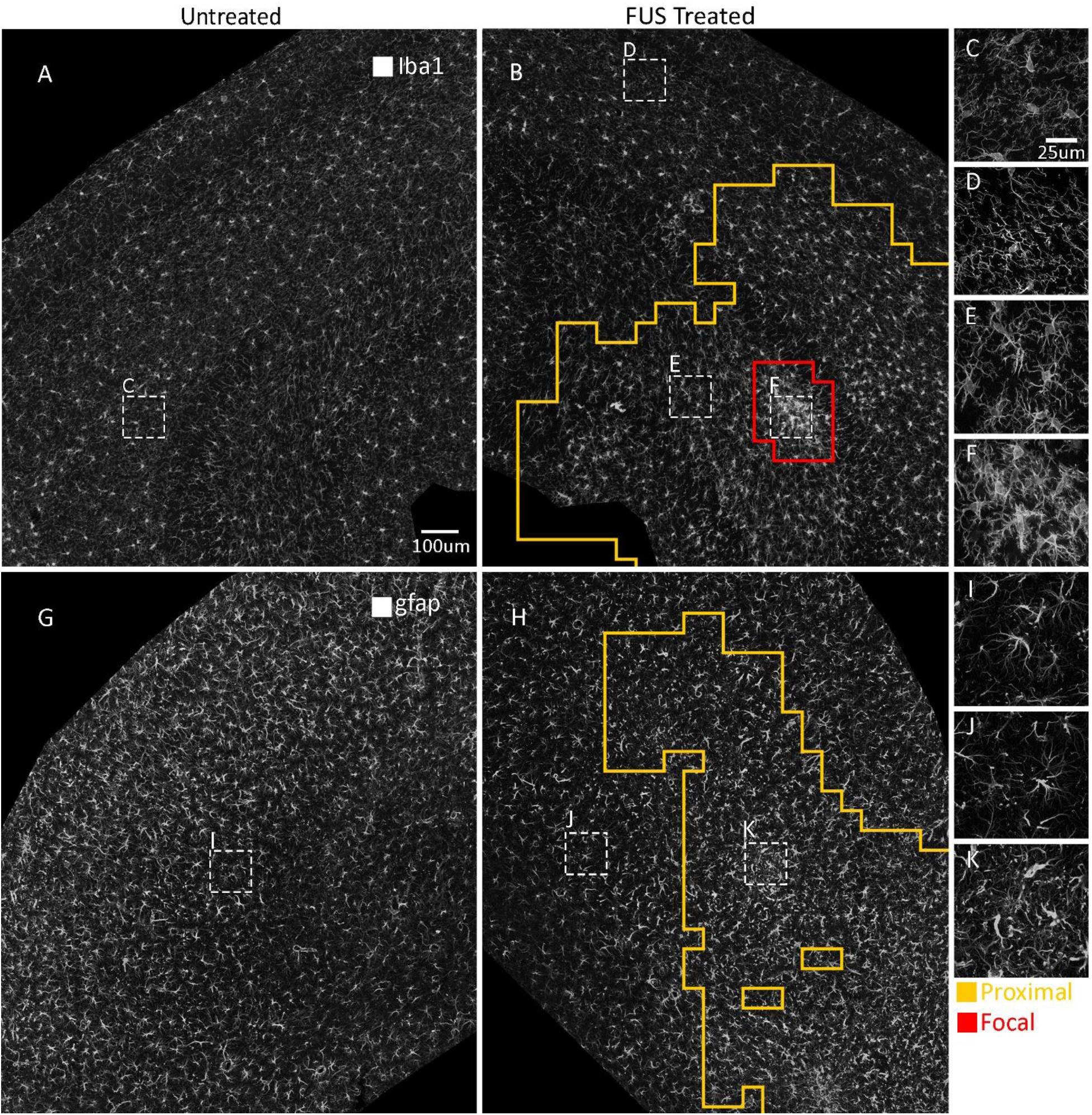
MORPHIOUS identifies regions of microglia and astrocyte activation following FUS treatment. A representative staining of Iba1+ microglia in contralateral (A) and FUS treated hippocampi (B) at 20x magnification. MORPHIOUS classified two regions of activation, proximal microglia (B, orange line) and focal microglia (B, red line). At high magnification (63x), contralateral microglia (C) as well as non-activated distal microglia (D), show a highly ramified morphology. (E) Proximal microglia show some de-ramification. (F) Focal microglia show substantial deramification and enlarged somas. A representative staining of Gfap+ astrocytes in contralateral (G) and FUS treated hippocampi (H) at 20x magnification. MORPHIOUS classified a single class of activated astrocytes, proximal astrocytes (H, orange line). At high magnification (63x), compared to contralateral (I) and distal (J) astrocytes, proximal astrocytes (K) show increased branching, and hypertrophy.

**Fig. 3.**
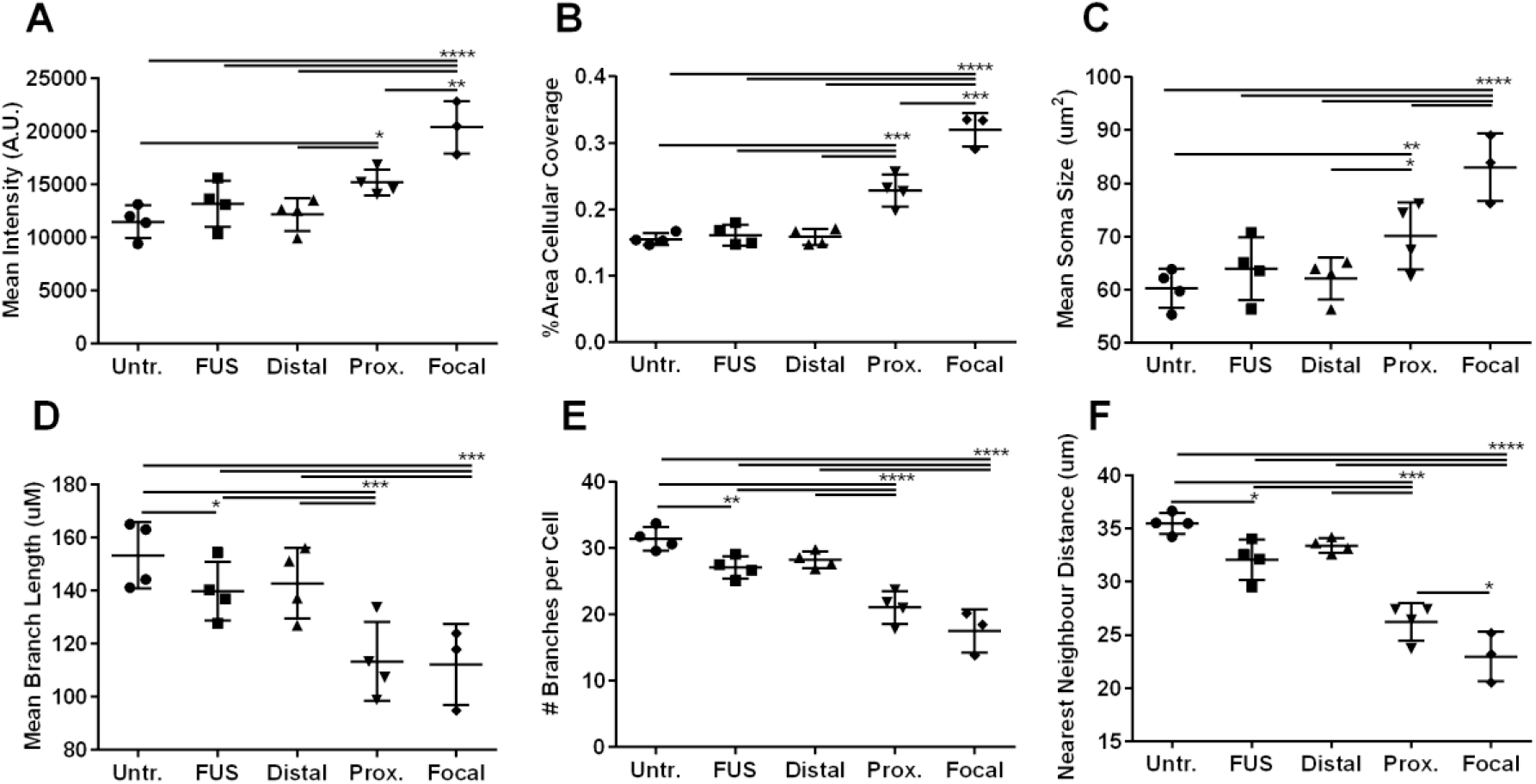
Focal and proximal microglia show morphological changes consistent with activation. The morphologies of untreated (Untr) microglia from contralateral hippocampi, FUS treated hippocampi (FUS), and MORPHIOUS classified distal, proximal (Prox.) and focal regions were compared. Cellular metrics included (A) Iba1 mean intensity, (B) Iba1 percent area coverage, (C) mean soma size, (D) mean branch length per cell, (E) number of branches per cell (F), and the nearest neighbour distance (G). Groups were analysed via a mixed linear model. Between-group differences were assessed via a Sidak’s post-hoc test. Significance: *P < 0.05; **P < 0.01; ***P < 0.001; ****P< 0.0001. Data represent means ± SD; n = 4 per group (Untr, FUS, Distal, Proximal) and n=3 per group (Focal).

It is conventional within biological analyses to evaluate gross histological changes by comparing the entirety of treated hippocampi to untreated examples. When comparing the entire FUS treated hippocampi to their contralateral side, whole FUS hippocampal sections showed a minor reduction in the branch length (P<0.01, Fig. 3D), number of branches (P<0.05, Fig. 3E), and nearest neighbour distance (P<0.05, Fig. 3F), but showed no changes for other features (P>0.05). Thus, MORPHIOUS could successfully isolate morphologically distinct groups of microglia which are not necessarily apparent from whole-section observations.

To further validate the activated state of focal and proximal microglia, we assessed microglia activation independently by co-staining Iba1 with CD68 and Tgf1β. CD68 has traditionally been used as a marker of both pro-inflammation^22–24^, and general microglia activation^25^, and has been shown to be expressed by microglia following FUS^17,26^. Tgf1β is considered to be an anti-inflammatory microglia marker and can facilitate neuroprotection^27^. Interestingly, in accordance with a gradient of activation, focal (vs. untreated: P<0.0001; vs. proximal: P<0.01) and proximal microglia (vs. untreated: P<0.01) showed progressively greater colocalization with Tgf1β (Fig. 5). Moreover, focal, but not proximal microglia colocalized with CD68 (vs. untreated: P<0.001).

#### Classifying Astrocytes

In response to FUS, MORPHIOUS classified a single class of activated astrocytes, which we termed proximal astrocytes. Compared to untreated contralateral astrocytes, proximal astrocytes exhibited a 1.3-fold increased Gfap intensity (P<0.001, Fig. 4A), a 1.5-fold increased area coverage (P<0.001, Fig. 4B), a 1.4-fold increased branch length (P<0.0001), and a 1.3-fold increased number of branches (P<0.05, Fig. 4C). As well, proximal astrocytes did not show changes to NND (P>0.05, Supplementary Fig. 5), which is consistent with findings demonstrating that *in vivo*, astrocytes do not migrate^20^. When comparing whole-contralateral and ipsilateral hippocampal slices, no features were significantly different. To validate MORPHIOUS predicted clusters of astrocyte activation, we observed that activated astrocytes colocalized with Nestin (vs. untreated, P<0.05), an intermediate filament protein which becomes upregulated during astrogliosis (Fig. 6). Moreover, there was a strong spatial overlap between activated astrocytes and microglia. In total, 15.8% and 10.3% of ipsilateral hippocampal sections were covered by activated microglia and astrocyte clusters, respectively (Supplementary Fig. 6C). Of this area, 75% of activated astrocytes overlapped with activated microglia, while 49% of activated microglia clusters overlapped with activated astrocyte clusters. Moreover, proximal cluster sizes for activated astrocytes strongly correlated with total cluster sizes for activated microglia (R^2^=0.753, P<0.0001, Supplementary Figure 6D). Collectively, this suggests that both cells are responding to the common FUS stimulus, and it provides evidence that both cell types are indeed activated.

**Fig. 4.**
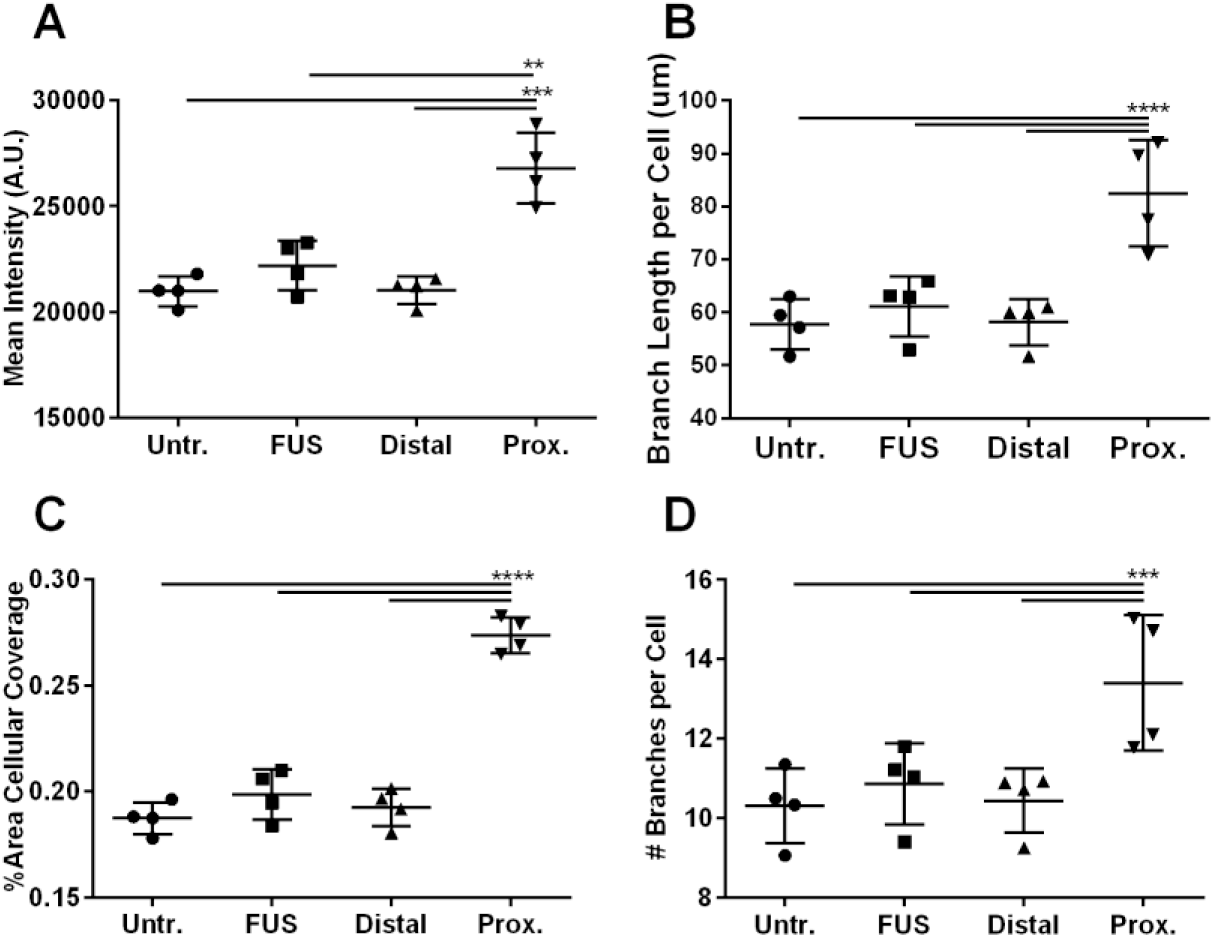
Proximal astrocytes show morphological changes consistent with activation. The morphologies of untreated (Untr) astrocytes from contralateral hippocampi, FUS treated hippocampi (FUS), and MORPHIOUS classified distal, and proximal (Prox.) regions were compared. Between-group differences in (A) Gfap mean intensity, (B) Gfap percent area coverage, (C) mean branch length per cell, and (D) mean number of branches per cell were assessed. Groups were analysed via a mixed linear model, and between-group were assessed via a Sidak’s Post-hoc analysis. Significance: *P < 0.05; **P < 0.01; ***P < 0.001; ****P< 0.0001. Data represent means ± SD; n = 4 per group (Untr, FUS, Distal, Proximal).

**Fig. 5.**
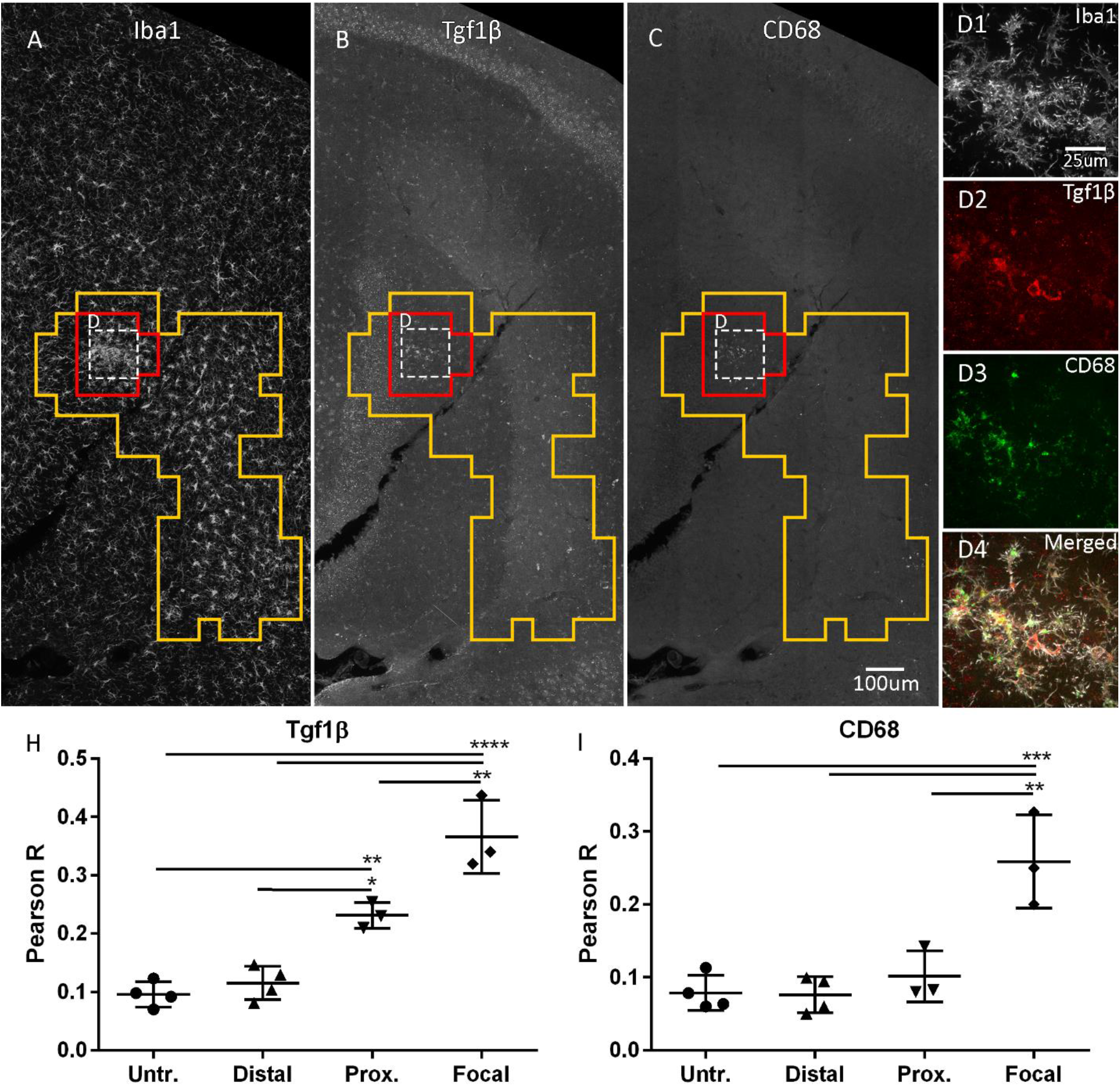
Focally activated microglia, but not proximally activated microglia upregulate common markers of activation. Focal (red line) microglia (A) colocalized with Tgf1b (B, D), and CD68 (C, D). Proximal (orange line) microglia colocalized with Tgf1b, but not CD68. Pearson correlation was used to colocalize Iba1 with Tgf1β (H) and CD68 (I). Images (A-C) were taken at 20x magnification. Insets (D1-D4) were taken at 63x magnification. Groups were analysed via a mixed linear model, and between-group were assessed via a Sidak’s Post-hoc analysis. Significance: *P < 0.05; **P < 0.01; ***P < 0.001; ****P< 0.0001. Data represent means ± SD; n = 4 per group (Untr, Distal, Proximal) or 3 per group (Focal).

**Fig. 6.**
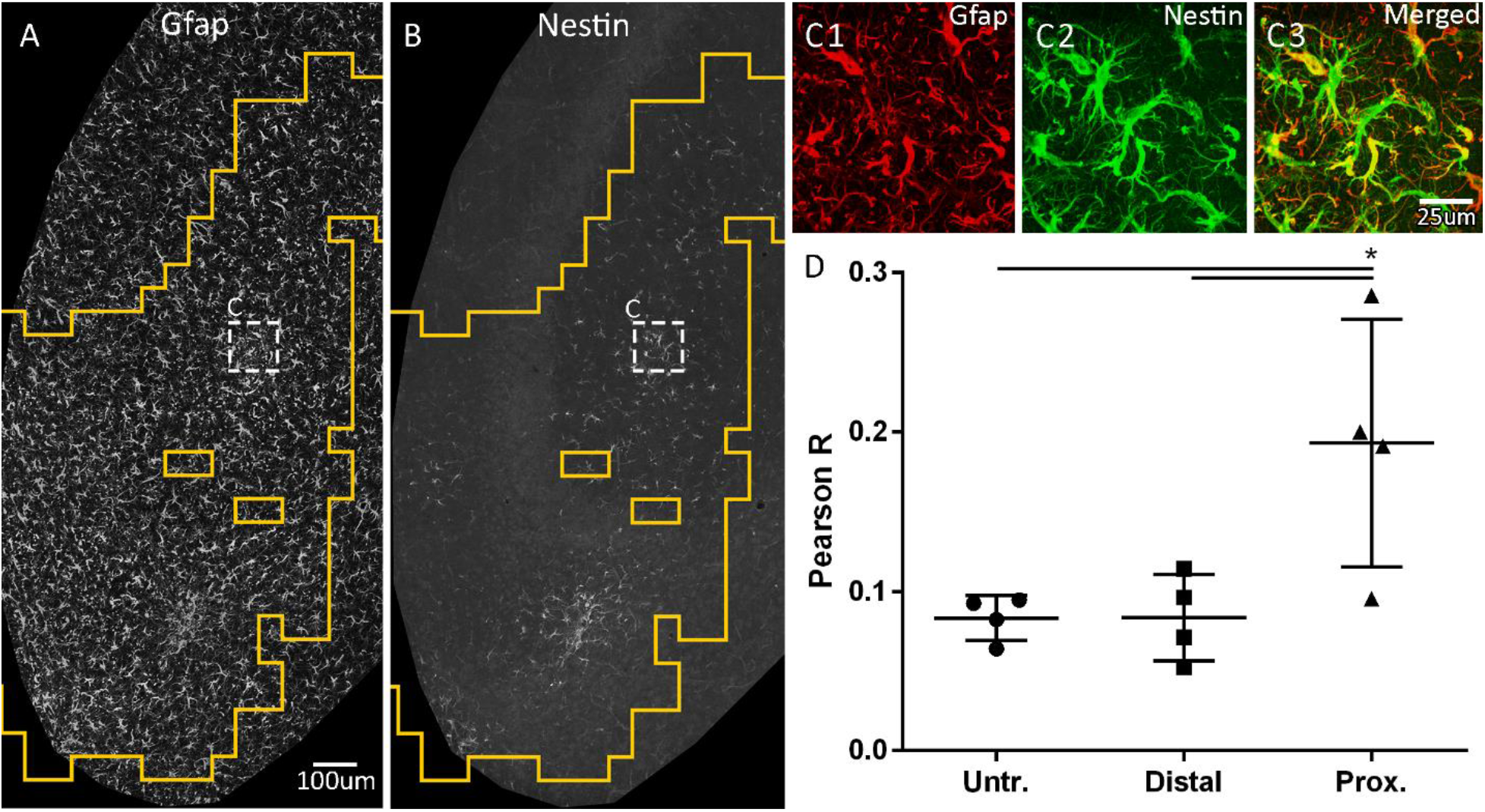
Proximal astrocytes co-express Nestin, a marker of activation. MORPHIOUS identified proximally activated (orange line) astrocytes (A) co-labelled with Nestin (B, C). Pearson correlation was used to colocalize Gfap with Nestin (D). Images (A-B) were taken at 20x magnification. Insets (C1-C3) were taken at 63x magnification. Groups were analysed via a mixed linear model, and between-group were assessed via a Sidak’s Post-hoc analysis. Significance: *P < 0.05; **P < 0.01; ***P < 0.001; ****P< 0.0001. Data represent means ± SD; n = 4 per group.

## Discussion

The unsupervised machine learning workflow MORPHIOUS distinguished heterogeneous regions of activated microglia and astrocytes, from surrounding non-activated tissues, while only referencing the morphologies of untreated glia. This capacity for the consistent identification and segmentation of activated clusters of glia, without the need for labelled examples of activation, promises to significantly improve the study glial activation in the brain, such as in response to disease progression or following treatment.

Following permeabilization of the BBB in the hippocampus using FUS, we identified two classes of activated microglia, focal microglia and proximal microglia which showed a gradient of activation features. Focal microglia, and to a lesser degree proximal microglia, showed elevated expression of Iba1, increased soma size, decreased nearest neighbour distance and decreased branching, features which are consistent with de-ramification^21^. At the molecular level, activated microglia can be identified by the expression of CD68^28^ and Tgf1β^29^, among other markers. In this respect, focal microglia colocalized with both CD68 and Tgf1β, providing additional evidence to support their classification as activated cells. Proximal microglia only colocalized with Tgf1β, and the degree of colocalization was less than that of focal microglia. Thus, focal and proximal microglia are consistent with gradations of activated microglia. In the same regions as activated microglia, MORPHIOUS independently identified clusters of astrocytes which are characterized by increased Gfap intensity, area coverage and branching. Increased branching and expression of Gfap are characteristic of astrogliosis^2^. Moreover, proximal astrocytes were colocalized with Nestin, an intermediate filament which colocalizes with Gfap when astrocytes are activated^30^. Thus, in addition to microglia, MORPHIOUS identified regions of morphologically distinct astrocytes which are consistent with glial activation.

Both microglia and astrocytes exhibit a range of activation phenotypes that cannot easily be encapsulated by traditional activation markers alone^31^. Indeed, single-cell RNA-sequencing analysis revealed that in response to a variety of environments, microglia show incredible complexity and heterogeneity with respect to their activation states, which is not accurately captured by previous definitions of activation (i.e., M1 and M2)^31,32^. Likewise genomic analysis of astrocytes has revealed significant heterogeneity in activation expression that is highly context specific^29^. Thus, by using a naïve analysis of morphological changes, MORPHIOUS is conducive towards a broader view of activation and can identify putative activation states of glia based on morphology alone. This approach also eliminates the need to co-stain for secondary activation markers and allows for the discovery and study of new activation states, based on changes in morphology alone. Moreover, by maintaining a naïve approach, MORPHIOUS reduces the need for limited and heterogeneous pathological, treated, and experimental tissue by inferring activation only through the referencing of readily available examples of healthy control tissue.

Our approach has significant advantages over previous work that identify activated glia through clustering in feature space alone, such as through k-means or hierarchical clustering^3–7^. While previous methods can evaluate the putative activation of individual cells, the heterogeneity in microglia morphology poses a risk for false positives, which are difficult to interpret given the nature of unlabelled data. For example, Davis et al.^3^ report that following orbital optic nerve crush, activated microglia were found distributed among resting microglia in both the treated and untreated olfactory bulbs, a surprising result which may be valid, but is difficult to evaluate without additional validation^3^. By contrast, MORPHIOUS avoids the inclusion of individual false-positive cells by clustering through a spatial approach. Compared to previous methods, MORPHIOUS is able to segment whole regions of activated glia; which is well suited for investigating the spatial nature of pathology in the brain and can be used to segment out regions of interest. For example, following stroke, previous characterization of microglia morphology within infarct and peri-infarct zones have relied either on qualitative definitions of these zones, or have used secondary immunohistochemical indicators^21^. By contrast, MORPHIOUS can identify heterogeneous clusters of glial activation, consistent with infarct and peri-infarct zones, based on *de novo* changes in glial morphology. In addition, MORPHIOUS may be adaptable to applications outside of identifying glial activation. Indeed, a similar one-class support vector machine approach has been used to segment the borders of tumors using MRI data ^33,34^.

Considering the fact that we do not have a ground truth for activated glia, we cannot rule out that we are over- or under-classifying the activation of glial cells. Tuning hyperparameters for one-class support vector machines is a critical, and often difficult task, for which a consensus on optimal methodology has yet to be reached^35^. A common technique is to maximize accuracy, while minimizing the number of false-positives, based on labeled data (i.e., the class of the data is known) ^33,34^. However, examples of positive-class cases (i.e. outlier data) can often be difficult to acquire. More advanced methods deploy a variety of strategies which focus on identifying patterns in the one-class itself in order to maximize the capacity to distinguish normal cases from outliers^35^.

In our case, we leveraged two plausible biological assumptions in order to optimize hyperparameters: 1) that activated glia will coalesce in spatial clusters that occur in response to a stimulus. This has been well documented to occur in cases of ruptured blood vessels^20,21^, or beta-amyloid plaques^16^. 2) that healthy control hippocampal brain tissue will exhibit no large clusters of outlier glia. Thus, in tuning our one-class support vector machine, we deployed a simple learning objective: find the set of hyperparameters which maximizes cluster size in the ipsilateral hippocampus while ensuring no clusters of activation are observed in untreated control hippocampal slices. Importantly, searching for clusters of outliers may not be suitable for images which are highly heterogenous, or, for identifying single, or small numbers of morphologically distinct cells. As with all machine learning approaches, the effectiveness of the learning model is limited by the range of features selected. To develop a simple and accessible approach, MORPHIOUS collects features through the widely used software ImageJ. The use of features with greater complexity, as recently described by other methods^4^, may further improve the range of classifications that can be identified. MORPHIOUS is adaptable to other unsupervised classifiers and could be further improved with state-of-the-art convolutional neural networks that can effectively interpret features from raw image files.

In conclusion, we demonstrate that MORPHIOUS can naïvely identify clusters of morphologically activated microglia and astrocytes. This novel unsupervised machine learning workflow was able to detect heterogenous discrete clusters of morphologically activated glia, as well as identify activated cells which did not expression common activation markers. Thus, MORPHIOUS is conducive to the naïve detection and novel discovery of activated glial cells, and establishes a new approach for the unsupervised classification of microglia and astrocytes.

## Material and Methods

### Animals

Male C57bl/6 mice (n=4) at 3.5 months of age were treated with focused ultrasound (FUS) unilaterally in the left hippocampus. Mice were sacrificed at 7 days (D) post-FUS and perfused with 4% paraformaldehyde, brains were extracted and post-fixed in 4% paraformaldehyde over night at 4°C. Brains were switched to 30% sucrose for >24 hours and sectioned at 40 μm using a microtome. Free floating sections were stored in cryoprotectant at −20°C until use. All procedures were conducted in accordance with guidelines established by the Canadian Council on Animal Care and protocols approved by the Sunnybrook Research Institute Animal Care Committee.

### Magnetic resonance imaging guided focused ultrasound

Prior to FUS treatment, mice were anesthetized with 5% isoflurane, and maintained at 2% isoflurane. Fur was removed from the head using depilatory cream. A 26-guage angiocatheter was inserted into the tail vein. Animals were imaged using a 7.0-T MRI (Bruker), and T2-weighted axial scans were used to target four focal spots targeting the hippocampus. FUS was conducted using an in-house system with a spherically focused transducer (1.68-MHz frequency, 75 mm diameter, 60 mm radius of curvature) and the BBB was permeabilized using standard parameters (10 ms bursts, 1 Hz burst repetition frequency, 120-s duration). At the initiation of sonication, mice were injected via the tail vein separately with Definity microbubbles (0.02 ml/kg; Lantheus Medical Imaging) and Gadovist (0.2 ml/kg, Schering AG). Each injection was followed by a 150ul flush with saline. Acoustic pressure was monitored using a using a polyvinylidene fluoride (PVDF) hydrophone coupled to a feedback controller. Acoustic pressure was increased incrementally after each pulse, until maximum pressure was detected, whereby pressure was reduced to and maintained at 25%. Maximum pressure was determined based on the detection of subharmonic emissions. BBB permeability was confirmed based on the presence of Gadovist enhancement on T1 weighted MR images.

### Immunofluorescence Staining

Serial sections (1:24) were antigen retrieved (10 mM sodium citrate, 80°C, 30 min), washed (phosphate buffered saline), blocked (5% Donkey Serum with 0.3% Triton-X), and incubated for 3 nights at 4°C with primary antibodies. Primary antibodies include rabbit anti-mouse Iba1 (Wako, cat: 016-20001; 1:500), goat anti-mouse Iba1 (Abcam, cat: ab107159; 1:1500), goat anti-Gfap (Santacruz Biotech, cat: sc-6170; 1:250), goat anti-mouse Gfap (Novus Biological, cat: NB100-53809; 1:2000), rabbit anti-Tgf1β (Abcam, cat: ab215715; 1:250), rat anti-CD68 (Biolegend, cat: 137002, 1:400), rat anti-ki67 (ThermoFisher, cat: 14-5698-82, 1:400), anti-rabbit s100β (Abcam, cat: ab41548; 1:1500). After washing, antibodies were incubated in secondary antibodies (Jackson Immunoresearch; 1:200) for 1 hour at room temperature, washed, and mounted.

### Imaging

Whole hippocampal slices were acquired using a Zeiss Z1 Observer/Yokogawa spinning disk (Carl Zeiss) microscope. Images were acquired using 40 μm z-stacks with a 1 μm step-size at with a 20X objective. All analysis was conducted using images at 20X magnification.

### Image processing and feature quantification

All image analyses procedures were performed using Fiji/ImageJ^36^.

#### Intensity and branching features

Microglia soma, branching, and intensity measures were visualized using Iba1 immunofluorescence. Similar to previous work, astrocytes were double labelled with S100b and Gfap^37^. S100b was used to demarcate soma, while branching and intensity measures were evaluated with Gfap. Z-stacked images were converted to maximum intensity projections. Prior to analysis, images were background subtracted, and despeckled. Images of astrocytes were contrast-enhanced. To collect features, for each image, a 100 μm x 100 μm sliding window was applied to the image which was iteratively translated across the image in the X and Y directions with a 50% overlap. For each iteration, a local threshold was applied (Method: Phansalkar, radius: 60, parameter 1: 0, parameter 2: 0). Immunoflourescence features (Mean, IntDen, Area) were quantified using the “Measure” command, and the fractal dimension (D) was measured using the “Fractal Dimension” command. Locally thresholded images were further binarized (i.e., “Binarize” command), skeletonized (i.e., “Skeletonize (2D/3D)” command) and branch features were collected (“Analyze 2D/3D Features”).

#### Cell soma features

Microglial and astrocytic cell bodies were segmented using custom scripts. Using custom python scripts, for each 100 x 100 μm window, mean soma area, soma circularity, and nearest neighbour distance (NND), were evaluated. Soma circularity was calculated using the formula: circularity = 4π(area/perimeter^2^). For each cell soma, the nearest neighbour distance was determined as the distance between the geometric center of a cell, and the nearest neighboring geometric cell center, as determined via Euclidean distance.

#### Segmenting microglia cells

To count microglia and astrocytes, we developed a custom macro using the MorphoLibJ library which segmented microglia and astrocyte cell bodies^19^. Iba1 images were first background subtracted by 50 pixels, and despeckled. Subsequently, using the MorphoLibJ library, we applied erosion (element: octagon, radius: 1), directional filtering (type: Max, operation: Mean, line: 6, direction: 32), morphological filter opening (element: Octagon, radius: 2), and top hat gray scale attribute filtering (attribute: Box Diagonal, minimum: 150, connectivity: 4). The resulting images were thresholded using the “IJ_IsoData” algorithm, and binarized. Cell body regions of interest (ROIs) were identified using the ImageJ particle analyzer command with a size filter of 25 pixels (scale: 1.5 pixels/μm).

#### Segmenting astrocyte cells

S100b images were first background substracted with a rolling ball radius of 50 pixels, and despeckled. Subsequently, using the MorphoLibJ library, we applied morphological filter opening (element: Octagon, radius: 2), gray scale attribute filter opening (attribute: Area, minimum:100, connectivity: 8), directional filtering (type: Max, operation: Mean, line: 10, direction: 32), and top hat gray scale attribute filtering (attribute: Box Diagonal, minimum: 100, connectivity: 4). A local threshold was applied to the resulting image (method: Phansalkar, radius: 60, parameter 1: −1, parameter 2: 0) which was subsequently binarized. Cell body ROIs were identified using the particle analyzer with a size filter of 30 pixels.

#### Feature selection and processing

Features used for identifying proximal microglia included area, mean intensity, the fractal dimension (D), number of cells, average NND, average soma size, average soma circularity, number of branches, branch length, number of branch junctions, number of triple branch points, number of branch ends, and the cellular perimeter. Features used for identifying proximal astrocytes included Area, Mean intensity, number of branch junctions, number of branch ends, number of slab branch pixels, number of triple points, and cellular perimeter area. Each feature was z-score normalized: z = (x - μ) / s, where x is the value, μ is the sample mean, and s is the sample standard deviation. Both contralateral and FUS-treated samples were normalized based on the mean and standard deviation of the contralateral dataset. Subsequently, features were transformed using principle component analysis, and enough principal components (PCs) were selected to retain 99% of variance. This corresponded to 9 PCs for the microglia feature set and 5 PCs for the astrocyte feature set. Z-score normalization and principal component analysis were conducted using the Scikit-learn module in python^38^.

### Identifying proximal clusters of microglia and astrocytes

A one-class support vector machine was trained using the features from contralateral sections. This model was used to identify outliers in FUS-treated sections. Since outliers can represent regions with either hyperintense features, or hypointense features, the initial set of putative outliers were filtered to ensure all identified outliers had a mean intensity that was larger than a z-score of −1. These candidate outliers were subsequently spatially clustered using the density-based spatial clustering of applications with noise (DBSCAN) algorithm ^39^. Outliers which subsequently spatially clustered were deemed proximal clusters. Implementations for the one-class support vector machine and DBSCAN were accessed from scikit-learn ^38^.

MORPHIOUS requires user input for four parameters: nu, gamma, minimum cluster size, and minimum neighbour distance. The nu and gamma parameters are hyperparameters for a one-class support vector machine, and the radial-basis-function kernel respectively. Nu reflects the percentage of normal observations which lie outside the classification decision boundary and is a regularization parameter. Gamma is a parameter for the radial basis kernel function. The minimum cluster size and distance are hyper parameters for DBSCAN which collectively defines the cluster size as the area where the number of points greater than the minimum cluster size are within the minimum neighbour distance. By default, MORPHIOUS sets the radius to be equal to the diagonal length of the window size rounded up (142 μm). Values for nu, gamma, and minimum cluster size for each stain were optimized via a grid search (Supplementary Fig. 2-3).

### Identifying focal clusters of microglia

We further classified a second subset of microglia, termed focal microglia. Conceptually, if proximal microglia are visualized as a feature-space hill, focal microglia represent the peak of this hill. To identify focal microglia, for each FUS-treated section, the mean Iba1 intensity of microglia are sorted in ascending order which results in an exponential curve (Fig. 1). The elbow point of this curve is subsequently used as a threshold value. Proximal windows with a mean Iba1 intensity which is greater than this threshold value are subsequently spatially clustered using DBSCAN, with a min cluster size of 5, and distance of 142 μm. To evaluate this elbow point, a vector is drawn to connect the first and last curve points (A_1_). Subsequently, a perpendicular vector B_x_ from every datapoint in the curve is connected to A_1_. The datapoint corresponding to the perpendicular vector with the greatest distance (i.e., max(|A_1_B_x_|)) was labelled as the elbow point. To ensure stability of elbow point, this procedure was iterated, and on each iteration, the first point in the curve was removed. As a result, the modal elbow point was used as the focal threshold value. Finally, to ensure that the proximal-mean Iba1 curve was sufficiently steep, in order to justify the existence of an exponential curve, distance |A_1_B_x_| must be greater than a threshold of 0.5, a value which worked well in our experience. Otherwise, focal clusters were not searched for.

### Parameter optimization

Using the contralateral datasets, 10-fold cross-validation was used to iteratively separate contralateral hippocampi into training and test sets, and identify the set of nu, gamma, and minimum cluster size parameters which resulted in no clustering within any contralateral hippocampi. A second grid search was run, training with the contralateral dataset, and testing on each FUS-treated hippocampus, to identify the set of hyper parameters which maximized cluster size within FUS-treated tissue. For each Iba1 (Supplementary Fig. 2) and Gfap antibody (Supplementary Fig. 3), a separate grid search was run, and optimal parameters identified.

### Colocalization analysis

Pearson correlation analysis was used to assess the colocalization between Iba1 and Tgf1β, Iba1 and CD68, and Gfap and Nestin. Colocalization analysis was conducted using the coloc2 plugin in ImageJ.

### Statistical analysis

Differences in cellular features were analyzed using a mixed-linear model. Main effect, pairwise between-group differences in cellular features were assessed with a Sidak post-hoc test. A value of p<0.05 was considered statistically significant. A linear regression was used to evaluate correlations between microglia and astrocyte cluster sizes, and between manual and automatic cell counts. All statistical analyses were conducted using SPSS (version 22, IBM).

### Source Code

The MORPHIOUS source code, as well as imageJ macros are available for use at https://github.com/jsilburt/Morphious.

## Acknowledgements

We thank Dr. Kullervo Hynynen for his expertise and assistance with using magnetic resonance imaging guided focused ultrasound (MRIgFUS). We acknowledge Shawna Rideout-Gros, Viva Chan, and Melissa Theodore for animal care, and Kristina Mikloska and Kairavi Shah for assistance with MRIgFUS. Finally, the authors would like to thank Dr. Dale Schuurmans for his discussions regarding machine learning and evaluation of the present method. This research was undertaken, in part, thanks to funding from the Canada Research Chairs program (I.A. Canada Research Chair in Brain Repair and Regeneration, Tier 1). This work was supported by the Canadian Institutes of Health Research (FRNs 137064 and 166184 to I.A.). Additional funding was received from the FDC Foundation, the WB Family Foundation, Gerald and Carla Connor, and the Weston Brain Institute (TR130117 to I.A.). Funding for staff and MRI procedures was received from the National Institute of Biomedical Imaging and Bioengineering of the National Institute of Health (RO1-EB003268, awarded to K.H.), and the Canadian Institutes of Health Research (FDN 154272, awarded to K.H.). J.S. was supported by a Frederick Banting and Charles Best Canada Graduate Scholarship and Ontario Graduate School Scholarship.

## References

1. Michell-Robinson, M. a. et al. Roles of microglia in brain development, tissue maintenance and repair. Brain 138, 1138–1159 (2015).

2. Pekny, M. & Pekna, M. Astrocyte Reactivity and Reactive Astrogliosis: Costs and Benefits. Physiol. Rev. 94, 1077–1098 (2014).

3. Davis, B. M., Salinas-Navarro, M., Cordeiro, M. F., Moons, L. & De Groef, L. Characterizing microglia activation: a spatial statistics approach to maximize information extraction. Sci. Rep. 7, 1576 (2017).

4. Salamanca, L. et al. MIC‐MAC: An automated pipeline for high‐throughput characterization and classification of three‐dimensional microglia morphologies in mouse and human postmortem brain samples. Glia 67, glia.23623 (2019).

5. Lu, Y. et al. Quantitative profiling of microglia populations using harmonic co-clustering of arbor morphology measurements. in Proceedings - International Symposium on Biomedical Imaging 1360–1363 (2013). doi:10.1109/ISBI.2013.6556785

6. Verdonk, F. et al. Phenotypic clustering: A novel method for microglial morphology analysis. J. Neuroinflammation 13, 153 (2016).

7. Fernández-Arjona, M. del M., Grondona, J. M., Granados-Durán, P., Fernández-Llebrez, P. & López-Ávalos, M. D. Microglia Morphological Categorization in a Rat Model of Neuroinflammation by Hierarchical Cluster and Principal Components Analysis. Front. Cell. Neurosci. 11, 235 (2017).

8. Karperien, A., Ahammer, H. & Jelinek, H. F. Quantitating the subtleties of microglial morphology with fractal analysis. Frontiers in Cellular Neuroscience 7, 1–34 (2013).

9. Sofroniew, M. V & Vinters, H. V. Astrocytes: biology and pathology. Acta Neuropathol. 119, 7–35 (2010).

10. Komura, D. & Ishikawa, S. Machine Learning Methods for Histopathological Image Analysis. Computational and Structural Biotechnology Journal 16, 34–42 (2018).

11. Kyriazis, A. D. et al. An End-to-end System for Automatic Characterization of Iba1 Immunopositive Microglia in Whole Slide Imaging. Neuroinformatics 17, 373–389 (2019).

12. Rostam, H. M., Reynolds, P. M., Alexander, M. R., Gadegaard, N. & Ghaemmaghami, A. M. Image based Machine Learning for identification of macrophage subsets. Sci. Rep. 7, 1–11 (2017).

13. Bzdok, D., Krzywinski, M. & Altman, N. Machine learning: supervised methods. Nat. Methods 15, 5–6 (2018).

14. Noble, W. S. What is a support vector machine? Nature Biotechnology 24, 1565–1567 (2006).

15. Schölkopf, B., Platt, J. C., Shawe-Taylor, J., Smola, A. J. & Williamson, R. C. Estimating the support of a high-dimensional distribution. Neural Comput. 13, 1443–1471 (2001).

16. Jordão, J. F. et al. Amyloid-β plaque reduction, endogenous antibody delivery and glial activation by brain-targeted, transcranial focused ultrasound. Exp. Neurol. 248, 16–29 (2013).

17. Kovacs, Z. I. et al. MRI and histological evaluation of pulsed focused ultrasound and microbubbles treatment effects in the brain. Theranostics 8, 4837–4855 (2018).

18. Ito, D., Tanaka, K., Suzuki, S., Dembo, T. & Fukuuchi, Y. Enhanced Expression of Iba1, Ionized Calcium-Binding Adapter Molecule 1, After Transient Focal Cerebral Ischemia In Rat Brain. Stroke 32, 1208–1215 (2001).

19. Legland, D., Arganda-Carreras, I. & Andrey, P. MorphoLibJ: integrated library and plugins for mathematical morphology with ImageJ. Bioinformatics 32, btw413 (2016).

20. Bardehle, S. et al. Live imaging of astrocyte responses to acute injury reveals selective juxtavascular proliferation. Nat. Neurosci. 16, 580–6 (2013).

21. Morrison, H. W. & Filosa, J. A. A quantitative spatiotemporal analysis of microglia morphology during ischemic stroke and reperfusion. J. Neuroinflammation 10, 4 (2013).

22. Kobayashi, K. et al. Minocycline selectively inhibits M1 polarization of microglia. Cell Death Dis. 4, e525–e525 (2013).

23. Perego, C., Fumagalli, S. & De Simoni, M. G. Temporal pattern of expression and colocalization of microglia/macrophage phenotype markers following brain ischemic injury in mice. J. Neuroinflammation 8, 174 (2011).

24. Xu, N. et al. Spared Nerve Injury Increases the Expression of Microglia M1 Markers in the Prefrontal Cortex of Rats and Provokes Depression-Like Behaviors. Front. Neurosci. 11, 209 (2017).

25. Boddaert, J. et al. CD8 signaling in microglia/macrophage M1 polarization in a rat model of cerebral ischemia. PLoS One 13, (2018).

26. Leinenga, G. & Gotz, J. Scanning ultrasound removes amyloid- and restores memory in an Alzheimer’s disease mouse model. Sci. Transl. Med. 7, 278ra33–278ra33 (2015).

27. Hu, X. et al. Microglial and macrophage polarization—new prospects for brain repair. Nat. Rev. Neurol. 11, 56–64 (2015).

28. Fu, R., Shen, Q., Xu, P., Luo, J. J. & Tang, Y. Phagocytosis of microglia in the central nervous system diseases. Mol. Neurobiol. 49, 1422–1434 (2014).

29. Neumann, J. et al. Microglia cells protect neurons by direct engulfment of invading neutrophil granulocytes: A new mechanism of CNS immune privilege. J. Neurosci. 28, 5965–5975 (2008).

30. Lin, R. C. S., Matesic, D. F., Marvin, M., McKay, R. D. G. & Brüstle, O. Re-expression of the intermediate filament nestin in reactive astrocytes. Neurobiol. Dis. 2, 79–85 (1995).

31. Ransohoff, R. M. A polarizing question: Do M1 and M2 microglia exist. Nature Neuroscience 19, 987–991 (2016).

32. Hammond, T. R. et al. Single-Cell RNA Sequencing of Microglia throughout the Mouse Lifespan and in the Injured Brain Reveals Complex Cell-State Changes. Immunity 50, 253–271.e6 (2019).

33. Zhou, J., Chan, K. L., Chong, V. F. H. & Krishnan, S. M. Extraction of brain tumor from MR images using one-class support vector machine. in Annual International Conference of the IEEE Engineering in Medicine and Biology - Proceedings 7 VOLS, 6411–6414 (2005).

34. Zhang, J., Ma, K.-K., Er, M.-H., Chong, V. & Hwa Er, M. Tumor Segmentation from Magnetic Resonance Imaging by Learning via one-class support vector machine Tumor Segmentation from Magnetic Resonance Imaging by Learning via one-class support vector machine TUMOR SEGMENTATION FROM MAGNETIC RESONANCE IMAGING BY LEARNING VIA ONE-CLASS SUPPORT VECTOR MACHINE. (2004).

35. Wang, S., Liu, Q., Zhu, E., Porikli, F. & Yin, J. Hyperparameter selection of one-class support vector machine by self-adaptive data shifting. Pattern Recognit. 74, 198–211 (2017).

36. Schindelin, J. et al. Fiji: An open-source platform for biological-image analysis. Nature Methods 9, 676–682 (2012).

37. Grosche, A. et al. Versatile and Simple Approach to Determine Astrocyte Territories in Mouse Neocortex and Hippocampus. PLoS One 8, e69143 (2013).

38. Pedregosa Fabianpedregosa, F. et al. Scikit-learn: Machine Learning in Python Gaël Varoquaux Bertrand Thirion Vincent Dubourg Alexandre Passos PEDREGOSA, VAROQUAUX, GRAMFORT ET AL. Matthieu Perrot. Journal of Machine Learning Research 12, (2011).

39. Ester, M., Kriegel, H.-P., Sander, J. & Xu, X. A Density-Based Algorithm for Discovering Clusters in Large Spatial Databases with Noise. (1996).

