## Supplementary Figures for "MORPHIOUS: A Machine Learning Workflow to Naively Detect the Activation of Microglia and Astrocytes"

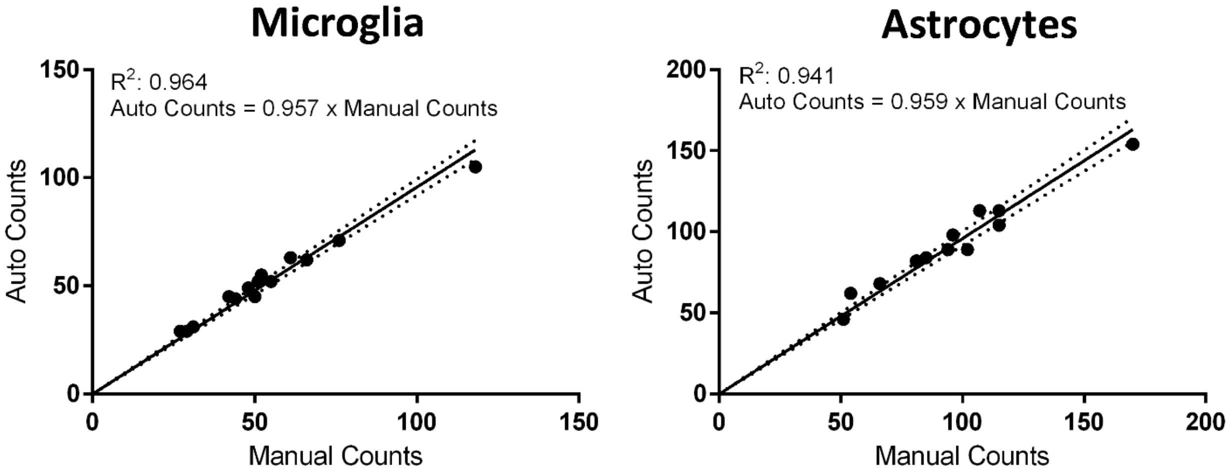


**Supplementary Fig. 1. Evaluating imageJ macros for the automated detection of glial cell bodies.**

(A) Microglia and (B) astrocyte cell bodies were detected from Iba1 and S100b immunofluorescence images using macros in imageJ. Using these protocols, predicted cell counts correlated strongly with manual cells counts, as evaluated using randomly selected fields of view. Comparisons were assessed using a linear regression model bounded through the origin. Significance: ****P< 0.0001.

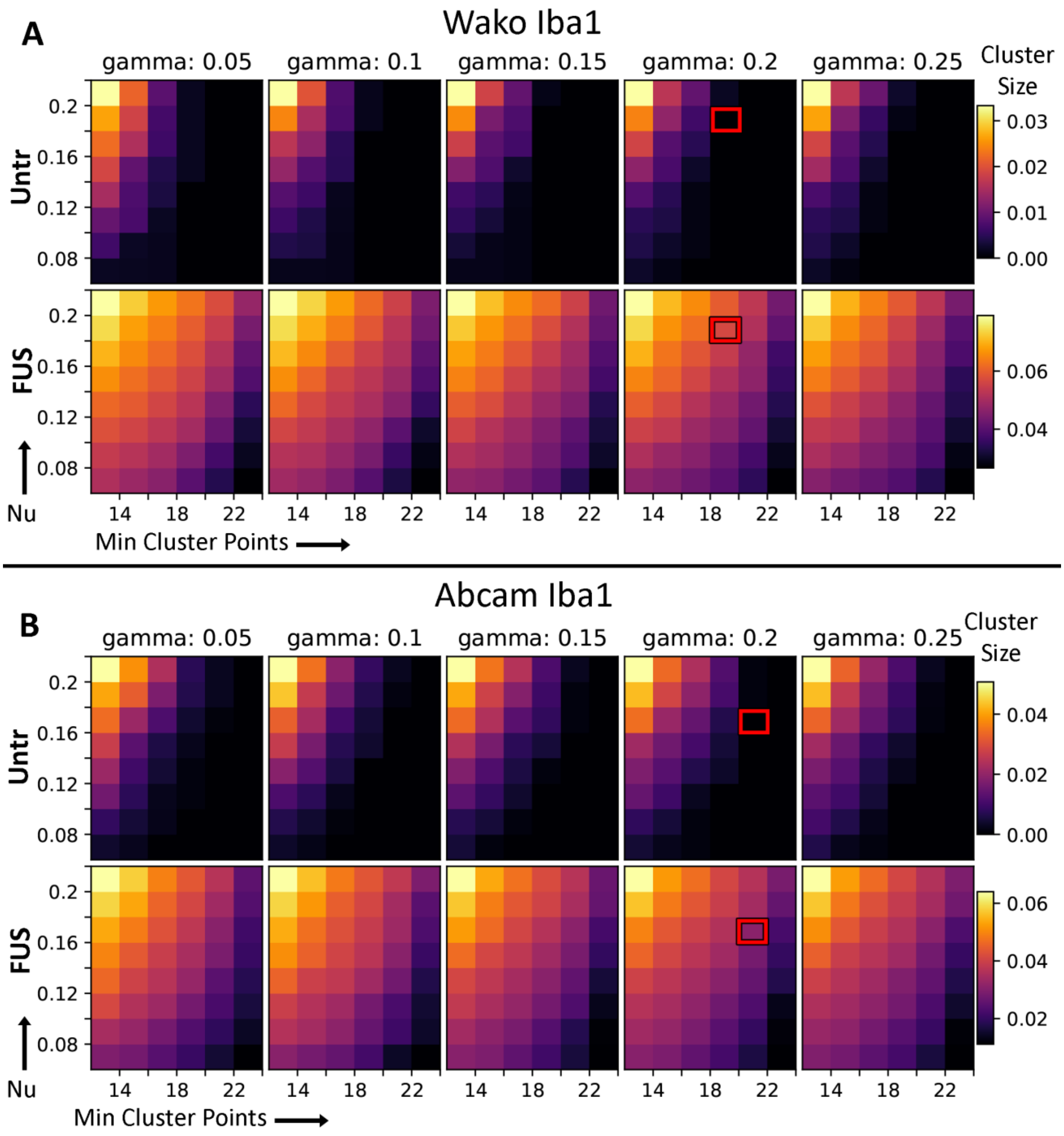


**Supplementary Fig. 2 Identifying MORPHIOUS hyperparameters for microglia activation clusters.**

Hyperparameters were determined for Wako (A) and Abcam (B) Iba1 antibodies. A grid search of Nu, Gamma, and minimum cluster points (min), were evaluated on the contralateral set of hippocampi, and the FUS treatment set using 10-fold cross validation. Final Hyperparameters (red rectangle) were selected as the set of hyperparameters which maximized the size of proximal microglia clusters in the FUS treatment dataset, while retaining no-false positive detections in the contralateral hippocampi. Final parameters were nu: 0.2, gamma: 0.2, min: 20 for the Wako Iba1 dataset, and nu: 0.2, gamma: 0.18, and min: 22 for the Abcam Iba1 dataset.

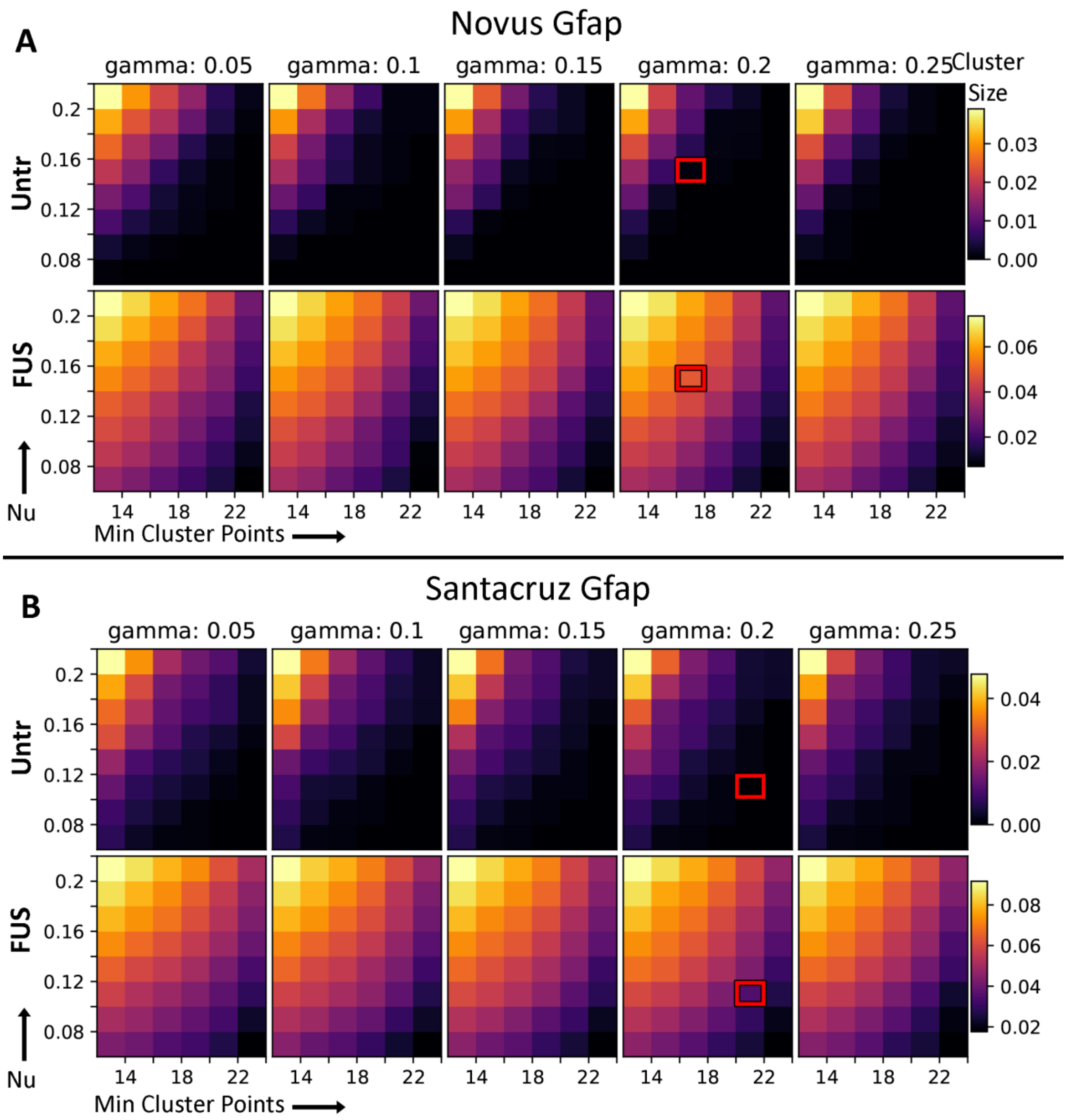


**Supplementary Fig. 3 Identifying MORPHIOUS hyperparameters for astrocyte activation clusters.**

Hyperparameters were determined for Novus (A) and Santacruz (B) Gfap antibodies. A grid search of Nu, Gamma, and minimum cluster size, were evaluated on the contralateral set of hippocampi, and the FUS treatment set using 10-fold cross validation. Final Hyperparameters (red rectangle) were selected as the set of hyperparameters which maximized the size of proximal microglia clusters in the FUS treatment dataset, while retaining no-false positive detections in the contralateral hippocampi. Final parameters were nu: 0.14, gamma: 0.2, min: 18 for the Novus Gfap dataset, and nu: 0.2, gamma: 0.12, and min: 22 for the Santacruz Gfap dataset.

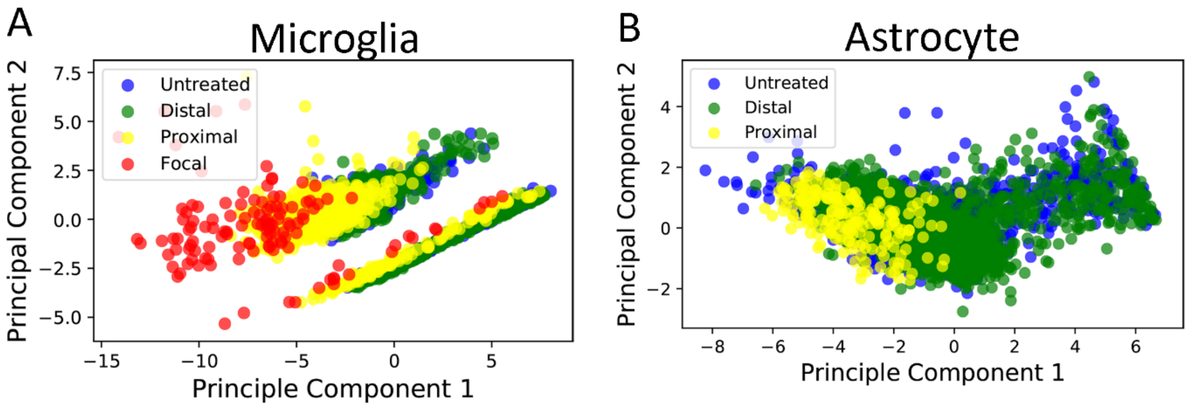


**Supplementary Fig. 4 Principle component analysis demonstrates that activated glia show distinct features.**

Principle component biplots of MORPHIOUS identified activated (A) microglia and (B) astrocytes demonstrate that focal and proximal microglia, and proximal astrocytes, occupy distinct locations in feature space. Non-activated, distal microglia and astrocytes, present in FUS treated hippocampi were indistinguishable from contralateral microglia and astrocytes, respectively. Principle component biplots are presented from representative sections for microglia and astrocytes, respectively.

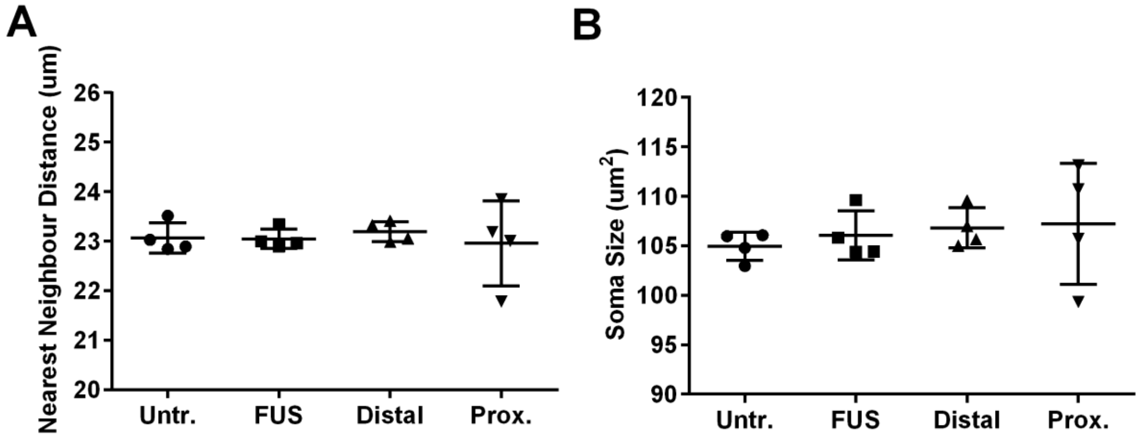


**Supplementary Fig. 5 Proximally activated astrocytes do not show changes in nearest neighbor distance or soma size.**

The morphologies of untreated (Untr) astrocytes from contralateral hippocampi, FUS treated hippocampi (FUS), and MORPHIOUS classified distal, and proximal (Prox.) regions were compared. Astrocyte soma were segmented using the astrocyte soma marker s100β, via custom ImageJ scripts. Between-group differences in (A) soma size, (B) nearest neighbour distance (NND) were evaluated. Groups were analysed via a mixed linear model, and between-group were assessed via a Sidak's Post-hoc analysis. Data represent means ± SD; n = 4 per group (Untr, FUS, Distal, Proximal).

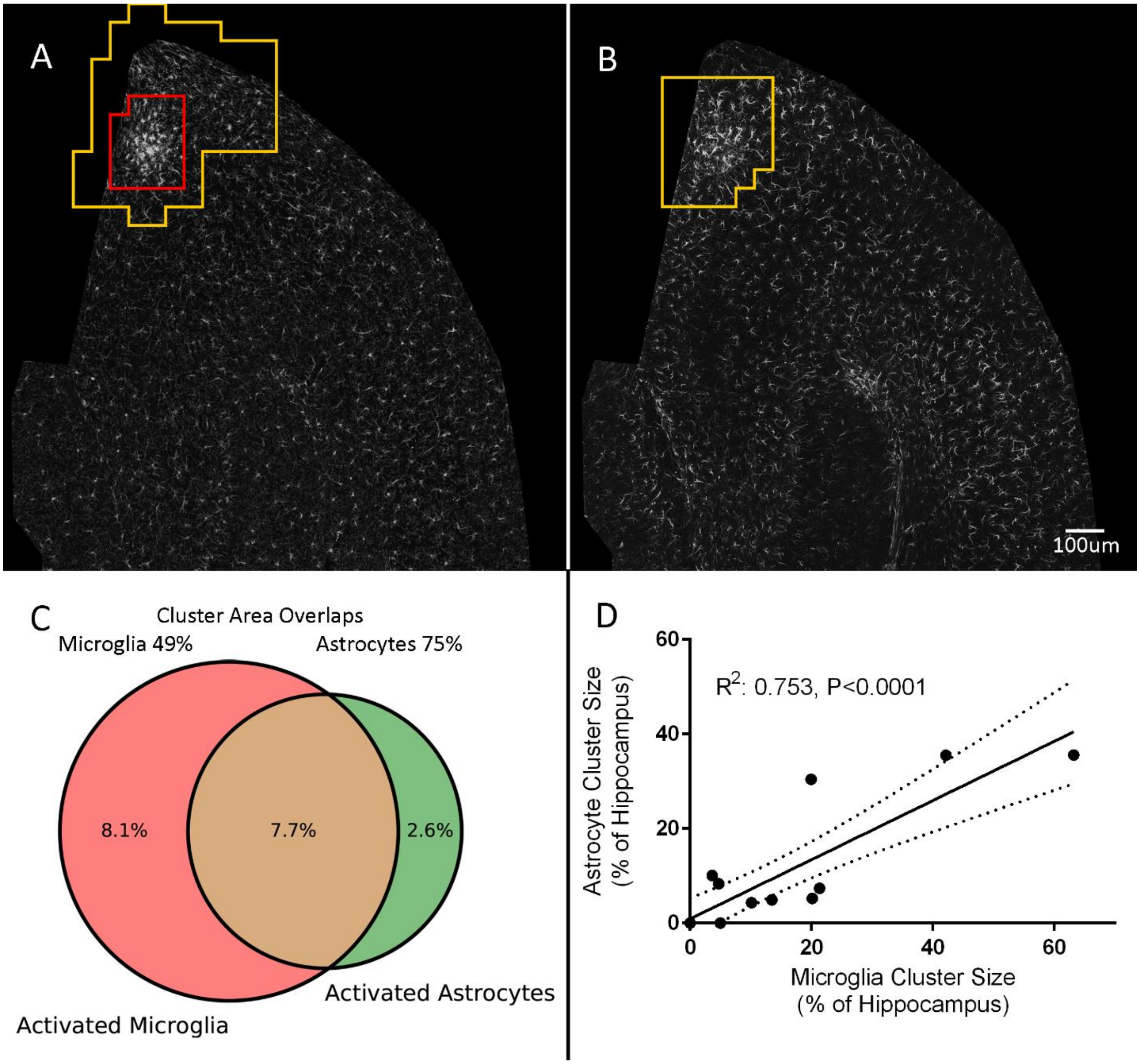


**Supplementary Fig. 6 Activated microglia overlap with activated astrocytes.**

Proximal clusters of microglia (A) overlap spatially with proximal clusters of astrocytes (B). Cluster sizes are reported as the percentage of the total hippocampal area covered by an activated microglia or astrocyte. In total, 74.5% of astrocyte clusters overlapped with microglia clusters, while 48.7% of microglia clusters overlapped with astrocytes. The correlation coefficient ($R^2$) was analyzed via linear regression analysis. Significance: ****P< 0.0001.
